# Predicting spectro-temporal modulation detection thresholds with a functional auditory model

**DOI:** 10.1101/2025.10.10.681587

**Authors:** Lily Cassandra Paulick, Helia Relaño-Iborra, Torsten Dau

## Abstract

Spectro-temporal modulation (STM) sensitivity has been proposed as a sensitive marker of speech intelligibility in challenging listening conditions, yet the underlying auditory mechanisms involved in STM detection remain incompletely understood. The present study measured STM detection thresholds in young normal-hearing and older hearing-impaired listeners and evaluated whether the revised Computational Auditory Signal Processing and Perception model [CASP, Paulick *et al*. (2025). J.Acoust.Am.**157**(5), 3232–3244] can account for individual performance. Thresholds were obtained for six modulation detection conditions, defined by combinations of spectral (0, 1, 2 c/o) and temporal (4, 12 Hz) rates. To individualise CASP, outer and inner hair cell loss estimates were obtained from audiometric and Adaptive Categorical Loudness Scaling (ACALOS) data. The results showed systematically elevated thresholds in older hearing-impaired listeners as compared to the young normal-hearing group, particularly at higher spectral rates. The model simulations reproduced overall threshold patterns, but substantially underestimated group differences and inter-individual variability in the data. Moreover, the simulations showed limited sensitivity to estimates of outer and inner hair cell loss, supporting the idea that additional supra-threshold mechanisms contribute to STM deficits. While these findings demonstrate the potential of auditory models to predict STM performance, they also highlighting the need for refined representations of peripheral and central processing to account for individual STM detection thresholds.

## I. INTRODUCTION

Accurately characterising the sources and consequences of individual hearing loss remains a central challenge in auditory science. A fundamental distinction is typically drawn between reduced audibility and supra-threshold deficits. Audibility limitations - commonly assessed with pure-tone audiometry or speech-in-quiet tests - reflect elevated detection thresholds and are often compensated for by amplification, restoring near-normal performance in quiet listening conditions in many cases (Plomp, 1978). Nevertheless, many listeners continue to experience difficulties when sounds are presented well above threshold, in so-called supra-threshold conditions (Dreschler and Plomp, 1985; Glasberg and Moore, 1989; Lopez-Poveda, 2014; Plomp, 1978). These difficulties are especially pronounced in noisy or complex environments and cannot be explained by audibility alone, pointing to additional deficits in auditory processing. Despite decades of research, the precise mechanisms underlying supra-threshold deficits remain only partly understood.

To investigate these mechanisms, numerous studies have examined supra-threshold deficits in specific auditory domains. Temporal processing has been assessed using tasks such as gap detection, temporal modulation transfer functions (TMTFs), and temporal fine structure sensitivity (Buss *et al*., 2004; Lorenzi *et al*., 2006; Oxenham and Moore, 1997; Qin and Oxenham, 2003; Regev *et al*., 2023). Spectral processing has been probed with measures of frequency selectivity and spectral masking (Dreschler and Plomp, 1985; Festen and Plomp, 1983; Glasberg and Moore, 1990; van Schijndel *et al*., 2001), while binaural processing has been evaluated through sensitivity to interaural time and level differences, as well as binaural masking level differences (Gabriel *et al*., 1992; Hall *et al*., 1984). A central motivation for these studies has been to determine whether such measures can predict individual speech-in-noise performance (Dreschler and Plomp, 1985; Houtgast and Festen, 2008; Johannesen *et al*., 2016; Strelcyk and Dau, 2009; Thorup *et al*., 2016). While informative, these measures typically explain only a modest portion of the variance in speech reception thresholds (SRTs) among hearing-impaired listeners.

Spectro-temporal modulation (STM) processing has received particular attention as a key ability supporting speech understanding (Singh and Theunissen, 2003; Venezia *et al*., 2019). STM detection thresholds measure sensitivity to combined temporal and spectral fluctuations, and, unlike other psychoacoustic tasks, have shown robust predictive power for speech intelligibility beyond the audiogram (Bernstein *et al*., 2016, Bernstein *et al*., 2013; Zaar/Simonsen and Laugesen, 2024). This has motivated the development of STM-based diagnostics, most prominently the clinically viable Audible Contrast Threshold (ACT) test (Zaar *et al*., 2023; Zaar/Simonsen and Laugesen, 2024). Early work by Chi *et al*. (1999) characterized STM detection in normal-hearing (NH) listeners, deriving spectro-temporal modulation transfer functions (MTFs) across a range of temporal (Hz) and spectral (cycles/octave, c/o) modulation rates. The MTFs showed low-pass characteristics in both dimensions, consistent with earlier findings on temporal resolution (Viemeister, 1979) and spectral resolution (Eddins and Bero, 2007; Green, 1986). Building on this, Bernstein *et al*. (2013) measured STM sensitivity in NH and hearing-impaired (HI) listeners using a four-octave pink noise carrier modulated at combinations of temporal (4, 12, 32 Hz) and spectral (0.5, 1, 2, 4 c/o) rates. HI listeners showed reduced sensitivity at low temporal and high spectral rates, with the largest group difference at 4 Hz and 2 c/o. Importantly, thresholds in this condition predicted speech intelligibility in stationary speech-shaped noise even after controlling for audibility (Bernstein *et al*., 2013). Follow-up studies using STM variants confirmed this link (Mehraei *et al*., 2014), leading to the development of the ACT test, which evaluates STM detection for a 4 Hz, 2 c/o condition imposed on a 2.5-octave-wide pink-noise carrier low-pass filtered at 2 kHz.

Despite its diagnostic potential, the auditory mechanisms underlying STM detection are not yet fully understood. Evidence suggests that STM performance reflects multiple processes, with temporal fine structure (TFS) encoding contributing more strongly at lower frequencies and frequency selectivity playing a greater role at higher frequencies (Bernstein *et al*., 2013; Mehraei *et al*., 2014). For example, Bernstein *et al*. (2013) reported that both low-rate (500 Hz) frequency modulation (FM) detection - linked to TFS sensitivity - and high-frequency (4 kHz) frequency selectivity - measured via notched-noise masking - jointly explained variance in STM performance (Bernstein *et al*., 2013; Summers *et al*., 2013). Extending this work, Mehraei *et al*. (2014) measured STM sensitivity using narrower one-octave carriers centred at 0.5, 1, 2, and 4 kHz. Their results showed that STM performance in HI listeners was frequency- and condition-specific, consistent with the involvement of distinct mechanisms. Importantly, after adjusting for audibility using the speech-intelligibility index (SII), STM thresholds at 4 Hz, 2 c/o for the 1 kHz carrier and 4 Hz, 4 c/o for the 4 kHz carrier remained significant predictors of speech intelligibility. Together, these findings support a dual-mechanism account in which TFS encoding at low frequencies and reduced frequency selectivity at higher frequencies jointly shape STM sensitivity.

Rather than relying solely on statistical associations between STM thresholds and individual psychoacoustic measures, the present study adopts a computational modelling approach using the Computational Auditory Signal Processing and Perception model (CASP, Jepsen and Dau, 2011; Jepsen *et al*., 2008; Paulick *et al*., 2025). Auditory models such as CASP provide a framework for identifying which auditory processing stages may limit performance in a given task and how hearing loss may alter these stages. A model that can predict individual STM thresholds has the potential to both help clarify the underlying mechanisms and to support the development of personalised diagnostic and compensatory strategies.

Earlier modelling work introduced the spectro-temporal modulation index (STMI), which combines a peripheral transformation with an explicit spectro-temporal analysis using a bank of modulation-selective filters tuned to different scales and rates. This model was consistent with TMTFs measured in NH listeners and successfully predicted speech intelligibility under various distortions (Chi *et al*., 1999; Elhilali *et al*., 2003). In contrast, CASP implements separate spectral and temporal modulation filterbanks, simulating nonlinear frequency selectivity and temporal modulation frequency selectivity, respectively. The present study evaluates whether CASP can account for STM detection thresholds without explicit spectro-temporal tuning. By doing so, we aim to determine the extent to which STM sensitivity can be explained by core auditory processing stages, as implemented in CASP, and to identify which aspects of hearing loss most strongly influence this ability.

The CASP model simulates auditory processing through a cascade of peripheral and central stages, including nonlinear frequency selectivity, adaptation mechanisms, and modulation frequency selectivity, followed by a decision stage based on an optimal detector designed for *n*-alternative forced choice (AFC) paradigms (Green and Swets, 1988). CASP has successfully predicted performance in a wide range of psychoacoustic tasks with NH listeners (Jepsen *et al*., 2008; Paulick *et al*., 2025) and has been extended to capture average effects of hearing loss (Jepsen and Dau, 2011). In addition, a speech-based variant of the model (sCASP, Relaño-Iborra *et al*., 2019) has been applied to predict speech intelligibility in noise.

In the present study, we measured STM detection thresholds in two listener groups: young normal-hearing (yNH) listeners and older hearing-impaired (oHI) listeners. We selected a subset of STM conditions from Bernstein *et al*. (2013), using temporal modulation rates of 4 and 12 Hz combined with spectral modulation rates of 1 and 2 c/o. For comparison, we also measured AM detection thresholds at the same temporal rates, but without spectral modulation. All modulations were imposed on a low-frequency pink noise carrier, consistent with the ACT paradigm (Zaar/Simonsen and Laugesen, 2024). To further characterise auditory processing in the oHI group, participants completed the Adaptive Categorical Loudness Scaling test (ACALOS, Brand and Hohmann, 2002), which provided individual estimates of nonlinear loudness growth. These parameters were incorporated into CASP to individualise the model’s front-end stages. Following established modelling frameworks (Chalupper and Fastl, 2002; Jürgens *et al*., 2011; Moore and Glasberg, 1997; Stefan *et al*., 2019), the overall audiometric hearing loss was decomposed into contributions from outer hair cell (OHC) and inner hair cell (IHC) dysfunction. This individualised modelling enabled us to simulate STM performance for each listener and to evaluate whether CASP can account for the observed across-listener variability.

## II. EXPERIMENTAL METHODS

### A. Listeners

Ten young normal-hearing (yNH) listeners (4 female, age range: 22-30 years, mean age = 23.9 years) and ten older hearing-impaired (oHI) listeners (2 female, age range: 54-80 years, mean age = 74.2 years) participated in the study. All listening tests were conducted monaurally. The oHI listeners had sensorineural hearing loss in both ears with no history of conductive impairment, and the tested ear was randomly selected. The yNH listeners had audiometric thresholds ≤ 20 dB HL at octave frequencies from 250 Hz to 8 kHz in the tested ear. Individual and mean audiograms of the tested ears are shown in Figure 1. All measurements were conducted without hearing-aid amplification. Participants provided written informed consent and were compensated for their time. The study protocol was approved by the Science Ethics Committee of the Capital Region of Denmark (Reference No. H-16036391).

**FIG. 1.**
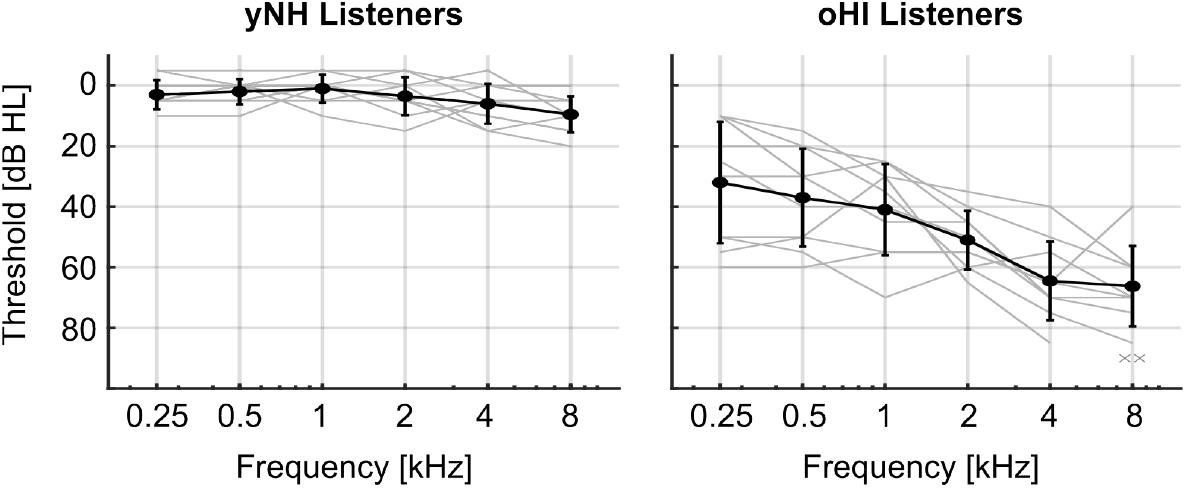
Audiometric thresholds for the tested ear of the young normal-hearing (yNH, left panel) and older hearing-impaired (oHI, right panel) listeners. Individual thresholds are shown in gray, with the mean and standard deviation across listeners shown in black.

### B. Procedure and Apparatus

All listening tests were conducted in single-walled, soundproof booths. Stimuli were generated in MATLAB at a sampling rate of 48 kHz, converted to analog via an RME Fireface soundcard (Haimhausen, Germany), and presented monaurally through Sennheiser HDA200 headphones (Wedemark, Germany). Output levels were calibrated using a GRAS RA0039 ear simulator (Holte, Denmark). NH listener completed one session in which they underwent pure-tone audiometry, otoscopy, and the AM and STM detection tasks. HI listeners completed two sessions in which they underwent the same tests, with the addition of the Adaptive Categorical Loudness Scaling (ACALOS) test. The test order was randomised across listeners, except for audiometry and otoscopy, which was always conducted at the beginning of the first session.

### C. (Spectro)-temporal modulation detection task

An STM stimulus consists of a carrier signal modulated by a combination of both spectral and temporal modulations, producing upward- or downward-moving ripples in the spectrum. The stimulus followed the design described by Zaar *et al*. (2023); Zaar/Simonsen and Laugesen (2024). The carrier was generated by summing 2499 random-phase tones, equally spaced along a logarithmic frequency axis spanning 2.5 octaves (354 to 2000 Hz), resulting in a pink-noise spectrum. STM thresholds were measured for four modulator conditions, defined by two spectral ripple rates (1 c/o and 2 c/o) combined with two temporal rates (−4 Hz and −12 Hz). Negative temporal rates correspond to upward-moving ripples. The modulator signal applied to the carrier was defined as

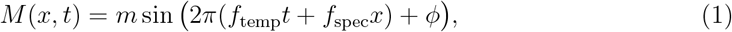

where *m* is the modulation depth, *f*_temp_ the temporal rate, *f*_spec_ the spectral rate, and *ϕ* the initial phase. A more detailed mathematical treatment can be found in Chi *et al*. (1999); Zaar *et al*. (2023). For comparison, pure temporal modulation thresholds (AM) were measured for the same two temporal rates. These stimuli were generated identically to the STM stimuli, except that the spectral modulation rate was set to zero.

STM thresholds were measured using a three-interval, three-alternative forced-choice (3I-3AFC) procedure with a 1-up-2-down tracking rule, targeting the 70.7% point on the psychometric function (Levitt, 1971). The procedure was implemented in MATLAB using the AFC-Toolbox 1.40 (Ewert, 2013). Each trial consisted of three intervals: two contained the unmodulated carrier and one contained the modulated carrier. The starting phase *ϕ* was randomly drawn from a uniform distribution in [−*π, π*] at the beginning of each run and held constant (‘frozen’) across all trials within that run. Each signal had a duration of 1 s, including 50 -ms raised-cosine onset and offset ramps, with 500 ms of silence between intervals. All stimuli were presented at a fixed level of 85 dB SPL. The adaptive variable was modulation depth, defined as *M* = 20 * log_10_(*m*) in dB, starting at *M* = −4 dB. The initial step size was 4 dB, reduced to 2 dB after the second reversal, and further reduced to a minimum of 1 dB. The threshold for each run was calculated as the mean modulation depth across the last six reversals at the minimum step size.

Thresholds were measured in three blocks (order randomised) defined by spectral modulation rate: pure AM (0 c/o), 1 c/o, and 2 c/o. Within each block, temporal rates of 4 and 12 Hz were tested in an interleaved manner, yielding six conditions in total. Fully modulated examples of these stimuli (M = 0 dB) are shown in Figure 2. Each condition was measured three times within the block, and the final threshold was computed as the mean across repetitions. Additional runs were performed if (i) a threshold was not reached, (ii) the standard deviation within a run exceeded 5 dB, or (iii) the standard error across repetitions exceeded 2 dB.

**FIG. 2.**
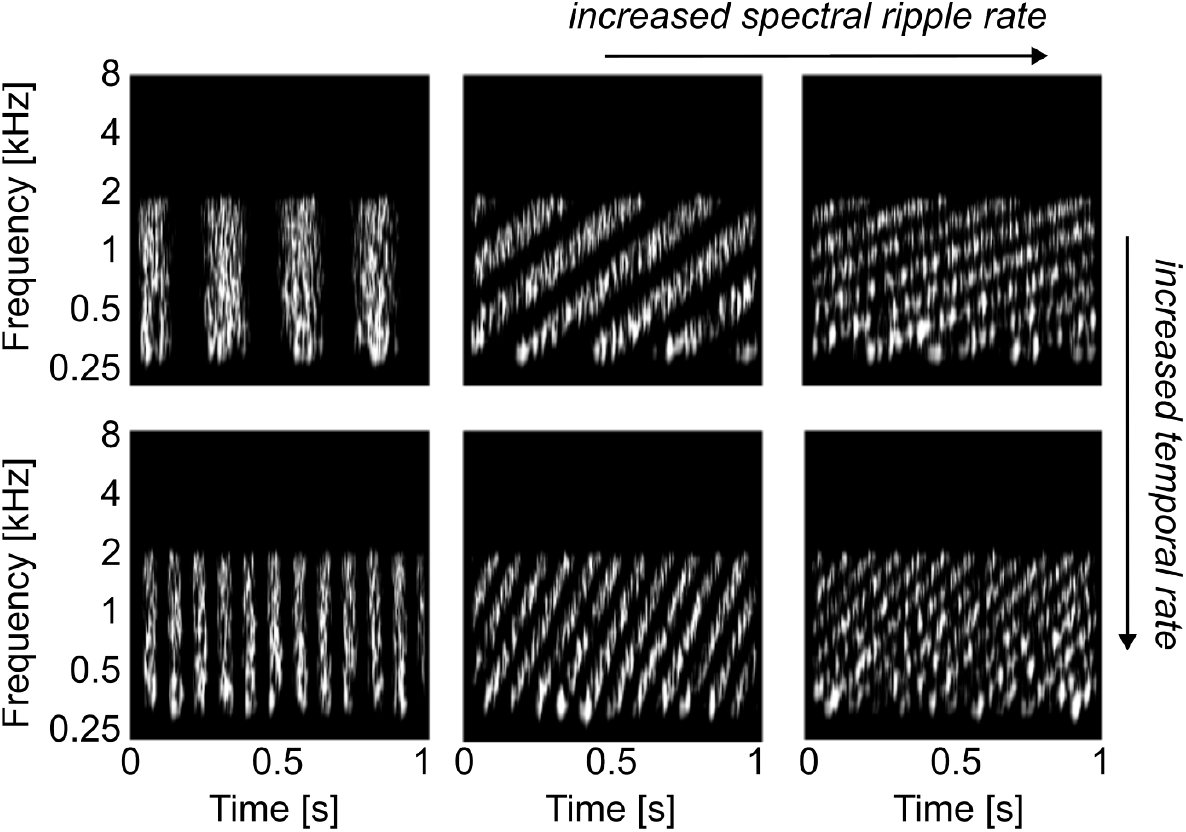
Auditory spectrograms of the stimuli used in the AM and STM detection tasks. Spectral ripple rates (0, 1 and 2 c/o; horizontal dimension) were combined with temporal modulation rates (4 and 12 Hz; vertical dimension). All modulations were applied to a 2.5-octave-wide noise carries with an upper cut-off frequency of 2 kHz. Spectrograms are shown for fully-modulated stimuli (M = 0 dB). During testing, modulation depth was adaptively varied to estimate the detection threshold for each AM and STM condition.

Participants received visual feedback after each trial indicating whether their response was correct. For STM conditions, one training run was completed per STM combination. For AM conditions, one training run was completed for the 4-Hz condition. If no threshold was reached in the initial training run, a second training run was provided.

### D. Adaptive categorical loudness scaling (ACALOS)

The ACALOS test (ISO 16832, 2006) was administered following the procedure described by Brand and Hohmann (2002). Stimuli were one-third-octave band noises with centre frequencies of 0.25, 0.5, 1, 2, 4, and 6 kHz. Each stimulus had a duration of 1 s with 50 ms raised-cosine onset and offset ramps. Participants rated the loudness of each stimulus on an 11-point categorical scale ranging from ‘inaudible’ to ‘extremely loud’. The maximum presentation level never exceeded 105 dB HL. Within each run, stimuli of different centre frequencies were presented in randomised, interleaved order. Each participants first completed a training run, followed by a second run from which the final results were obtained. For details on the ACALOS procedure the reader is referred to Brand and Hohmann (2002).

### E. Statistical Analysis

Statistical analyses were performed on the measured (spectro)-temporal modulation detection thresholds using a repeated-measures analysis of variance (ANOVA) based on fits of a linear mixed-effects model (LMM). The model included group (yNH versus oHI), tempo-ral modulation rate, and spectral modulation rate as fixed effects, and listener as a random effect. All main effects and their interactions were tested. Model assumptions were checked using Levene’s test for homogeneity of variance and the Shapiro-Wilks test for normality of residuals. Significant interactions were followed up with Holm-Bonferroni-corrected post hoc pairwise comparisons. The significance level was set to 0.05 for all analyses.

## III. AUDITORY PROCESSING MODEL

The preprocessing stages of the CASP model are illustrated in Figure 3 and follow the framework described by Jepsen *et al*. (2008); Paulick *et al*. (2025). Briefly, the input signal passes through outer- and middle-ear filters, followed by a dual-resonance nonlinear filter-bank (DRNL, Lopez-Poveda and Meddis, 2001) that simulates level-dependent frequency selectivity. The filterbank output is then processed by a nonlinear IHC stage, introduced by Paulick *et al*. (2025), which captures saturation at high sound pressure levels. This is followed by an adaptation stage consisting of five cascaded feedback loops (Püschel, 1988). Finally, the signal is decomposed into modulation sub-bands using a modulation filterbank, yielding a three-dimensional internal representation with dimensions of time, auditory frequency, and modulation frequency.

**FIG. 3.**
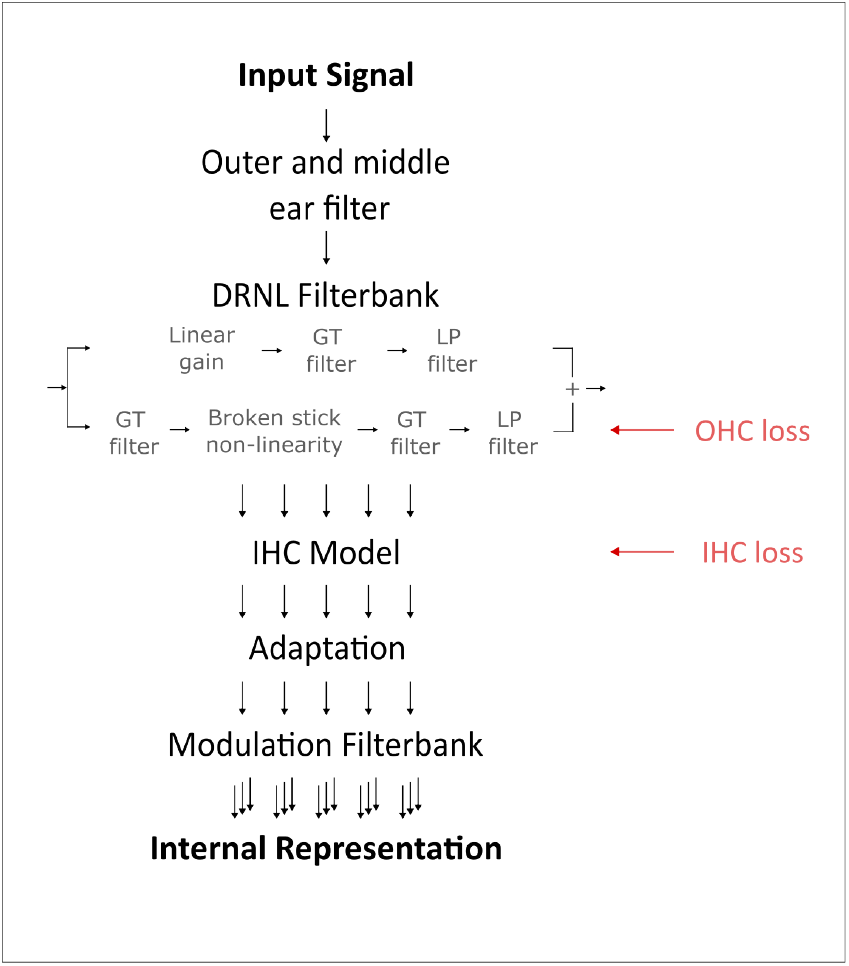
Preprocessing stages of the CASP model that construct the three-dimensional internal representation. Sensorineural hearing loss is simulated via (i) outer hair cell (OHC) loss, implemented as a modification of the broken-stick nonlinearity in the dual-resonance non-linear (DRNL) filterbank, and (ii) inner hair cell (IHC) loss, implemented as an attenuation at the output of the IHC stage.

The model backend consists of an optimal detector designed for *n*-AFC paradigms. The model follows the same adaptive procedure as the listeners and makes decisions on each trial based on correlations between a template and the interval representations. For the present modulation detection tasks, templates were generated with a modulation depth of *M* = 0 dB and averaged across 15 presentations, as in Paulick *et al*. (2025). Template generation was performed within-subject, under the assumption that each simulated HI listener forms an individual template. The model also incorporates internal noise that limits resolution. As in previous implementations, the variance of the internal noise was calibrated in an intensity-discrimination task to match just-noticeable differences (JNDs) at 60 dB SPL for both a 1-kHz tone and broadband noise (Paulick *et al*., 2025). This calibration was performed in the NH configuration and subsequently kept constant across tasks and simulated HI listeners.

Among the preprocessing stages, the DRNL filterbank plays a central role in simulating sensorineural hearing loss (SNHL). The DRNL filterbank implements level-dependent frequency selectivity using two parallel pathways: a linear path controlled by a gain parameter, and a nonlinear path incorporating a broken-stick nonlinearity. Together, these pathways shape the DRNL input-output (I/O) function:

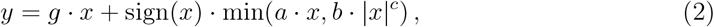

where *g* sets the slope of the linear region (dominating at higher levels), *c* is the compression exponent, and *a* and *b* determine the knee point where compression begins.

To simulate SNHL within this framework, it is assumed that the total audiometric hearing loss can be separated into contributions from OHC and IHC dysfunction (Jepsen and Dau, 2011; Lopez-Poveda and Johannesen, 2012; Moore and Glasberg, 1997):

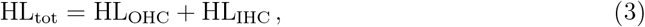

with both terms expressed in dB. This formulation assumes that audiometric thresholds reflect SNHL alone, excluding other sources of impairment. The OHC-related component, HL_OHC_, reflects a loss of cochlear gain, typically observed in basilar-membrane input–output (BMIO) functions or loudness growth functions. Earlier CASP work (Jepsen and Dau, 2011) estimated this parameter from BMIOs derived from temporal masking curves (Lopez-Poveda *et al*., 2003; Plack and Oxenham, 1998). In the present study, HL_OHC_ was instead estimated from ACALOS data using the procedure described by Jürgens *et al*. (2011). Briefly, individual loudness functions were derived from ACALOS and fitted with a dynamic loudness model (Chalupper and Fastl, 2002) that incorporates the two-component framework of hearing loss, separating OHC- and IHC-related contributions. The model simulates loudness growth for different proportions of OHC impairment, and a fitting algorithm determines the proportion that best matches the measured data. This approach yields time-efficient estimates consistent with those from TMCs (Jürgens *et al*., 2011).

Based on the estimated HL_OHC_, the DRNL I/O function (Equation 2) was then modified. Lowering *a* raises the compression knee point, reducing low-level gain, while lowering *b* extends compression to lower levels. Because many parameter combinations can yield similar I/O functions, OHC-driven modifications were restricted to the *a* parameter, keeping *b, c*, and *g* fixed:

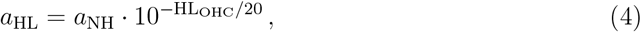

where *a*_NH_ is the normal-hearing value and *a*_HL_ the adjusted value for OHC loss. The residual hearing loss, after accounting for HL_OHC_, was attributed to IHC dysfunction, HL_IHC_, and implemented as an attenuation applied to the IHC stage outputs.

Select adjustments were made to the model implementation compared to that described in Paulick *et al*. (2025), specifically in the DRNL filterbank parametrisation, the modulation phase sensitivity implementation and backend metric. Details regarding these adjustments, as well as a backwards-compatibility assessment using the same benchmark experiments as in Paulick *et al*. (2025), are given in the Appendix.

## IV. RESULTS

### A. Experimental results

Modulation detection thresholds for the yNH listeners (blue) and the oHI listeners (red) are shown in Figure 4 as a function of spectral rate (0, 1, and 2 c/o) at two temporal rates: 4 Hz (left panel) and 12 Hz (right panel). The statistical analysis revealed significant main effects of group (*F*_1,18_ = 23.65, *p <* 0.001), temporal rate (*F*_1,90_ = 27.36, *p <* 0.001), and spectral rate (*F*_2,90_ = 163, *p <* 0.001). Significant two-way interactions were also observed between temporal and spectral rates (*F*_2,90_ = 12.18, *p <* 0.001) and between spectral rate and group (*F*_2,90_ = 41.9, *p <* 0.001). Neither the interaction of temporal rate and group nor the three-way interaction reached significance. Post-hoc analyses confirmed that oHI listeners had significantly higher (worse) thresholds than yNH listeners overall (*p <* 0.001). Group differences, expressed as the modulation depth difference Δ*M*, were especially pronounced at the highest spectral rate, i.e. at 2 c/o at 4 Hz (Δ*M* = 7.43 dB, SE = 0.99 dB) and 2 c/o at 12 Hz (Δ*M* = 5.62 dB, SE = 0.99 dB). These results indicate that both temporal and spectral rates significantly affect STM detection, and that oHI listeners are disproportionately affected by increases in spectral rates.

**FIG. 4.**
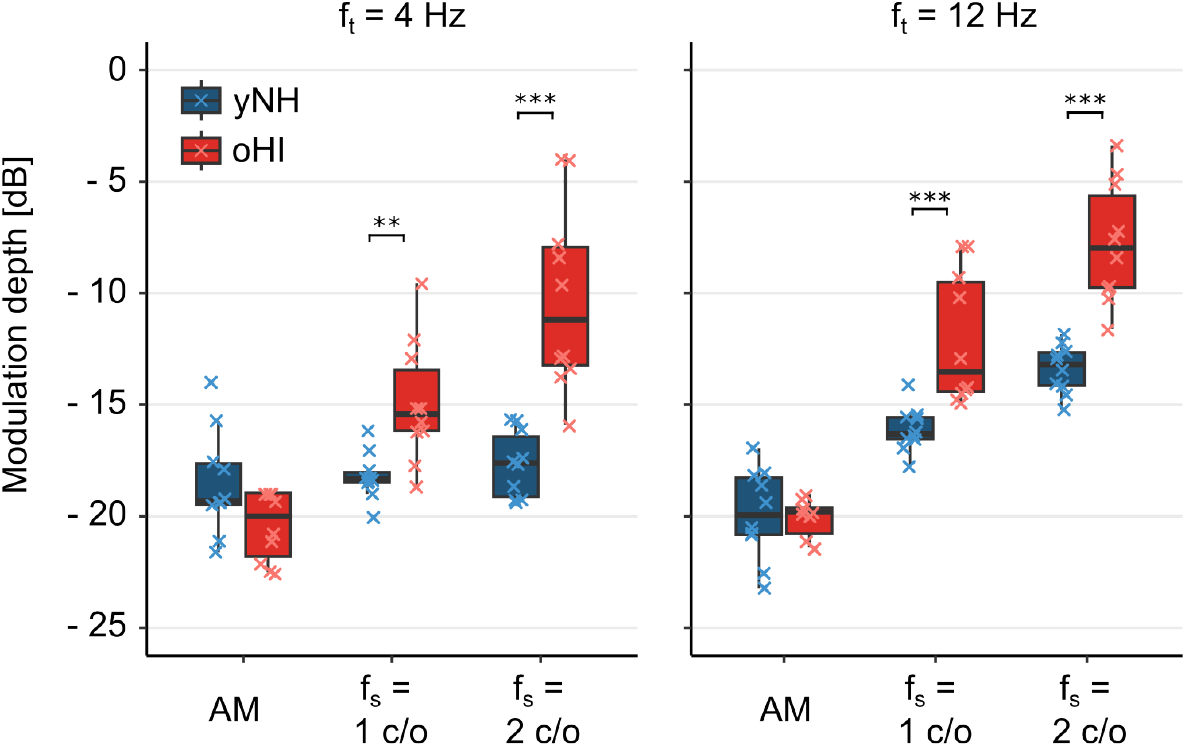
Boxplots of (spectro)-temporal modulation detection thresholds for yNH (blue) and oHI (red) listeners across temporal and spectral rate combinations. Left: Thresholds at 4 Hz as a function of spectral rate (0, 1 and 2 c/o). Right: Thresholds at 12 Hz. Individual data are shown as crosses. In boxplots, boxes represent the interquartile range (IQR; 25th to 75th percentiles), central lines indicate medians, and whiskers extend to 1.5 × IQR. Stars indicate the significance level of group differences (* *p* < 0.05, ** *p* < 0.01, *** *p* < 0.001).

### B. Model predictions

The top panels of Figure 5 show AM and STM thresholds for yNH listeners (dark-blue boxplots) alongside CASP predictions (light-blue circles). The model reproduced the general trend of lower thresholds at 4 Hz compared to 12 Hz, and the monotonic increase with spectral rate at 12 Hz. However, at 4 Hz, the model overestimated AM sensitivity and predicted a performance drop from AM to STM that was not observed in the data, thereby exaggerating the AM-STM difference. Quantitatively, predictions for NH listeners yielded a mean absolute error (MAE) of 2.18 dB and a strong correlation with empirical means across conditions (Pearson’s *r* = 0.89). The largest deviation (5.13 dB) occurred in the 4 Hz AM condition.

**FIG. 5.**
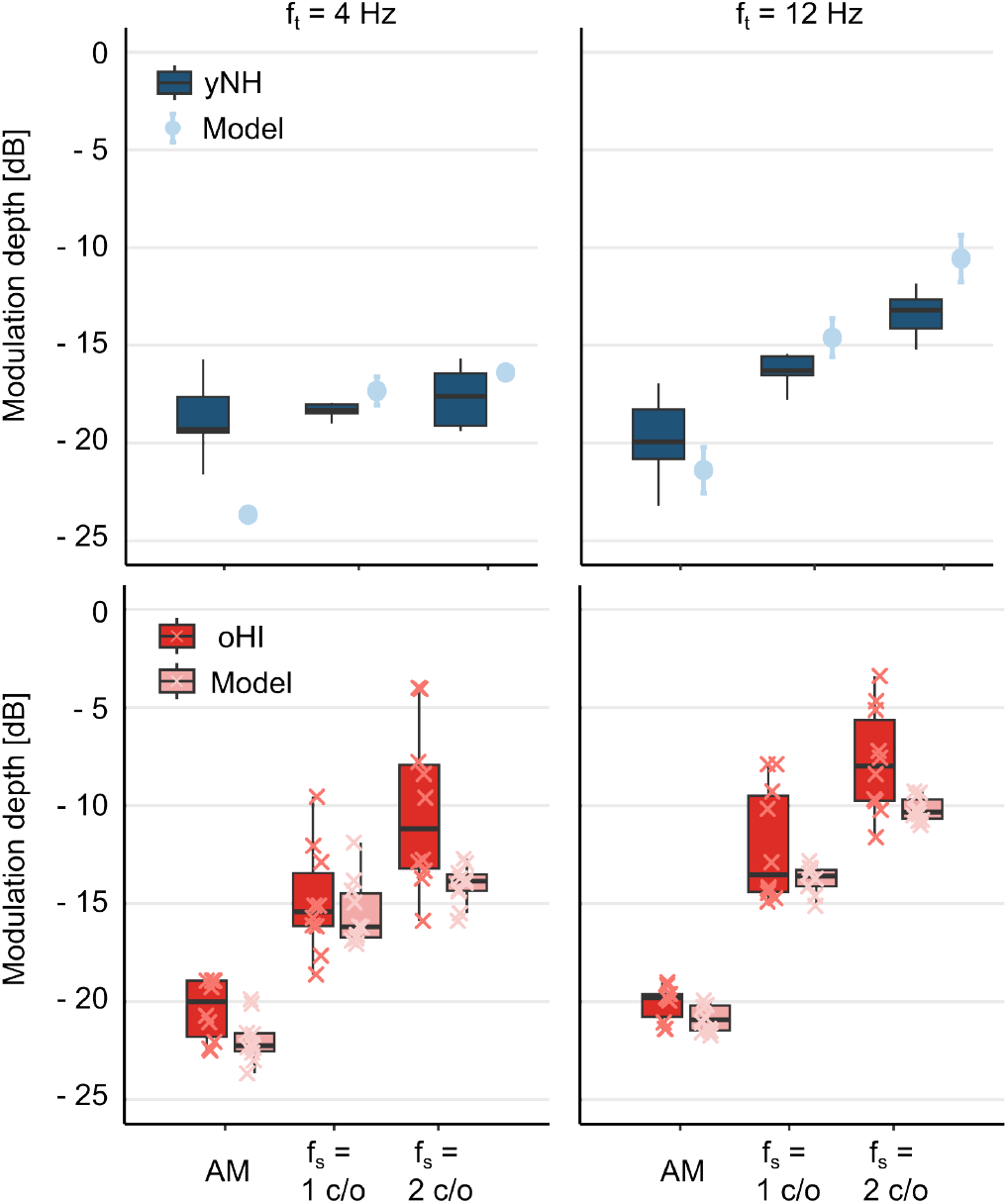
Top: AM and STM thresholds for yNH listeners (dark-blue boxplots) and CASP predictions (light-blue symbols with errorbars). Error bars represent the SD across repeated simulations. The model achieved a mean absolute error of 2.18 dB and correlated strongly with listener data (*r* = 0.89). Bottom: AM and STM thresholds for oHI listeners (dark-red, individual data shown as crosses) and CASP predictions (light-red, individual data shown as crosses). Across conditions, HI predictions yielded a MAE of 1.81 dB and a strong correlation with group averages (*r* = 0.98). For all boxplots, boxes represent IQRs, central lines medians, and whiskers extend to 1.5 × IQR.

The bottom panels of Figure 5 show measured and predicted thresholds for oHI listeners. Predictions relative to the group average yielded a MAE of 1.81 dB and a strong correlation with empirical means (*r* = 0.98). The largest deviation (3.82 dB) occurred in the 4 Hz, 2 c/o condition. Despite this close match on average, the model underestimated both the variability in the listener data and the magnitude of the group effect. This discrepancy is illustrated in Figure 6, which shows mean group differences in threshold (ΔM = M_HI_ −M_NH_) across conditions. Behaviourally, HI listeners performed similarly to NH listeners in AM detection but showed elevated thresholds in STM conditions, with group differences of up to ∼ 7.43 dB. In contrast, CASP predicted only modest threshold elevations (up to ∼ 2.33 dB), primarily in the 4 Hz conditions.

**FIG. 6.**
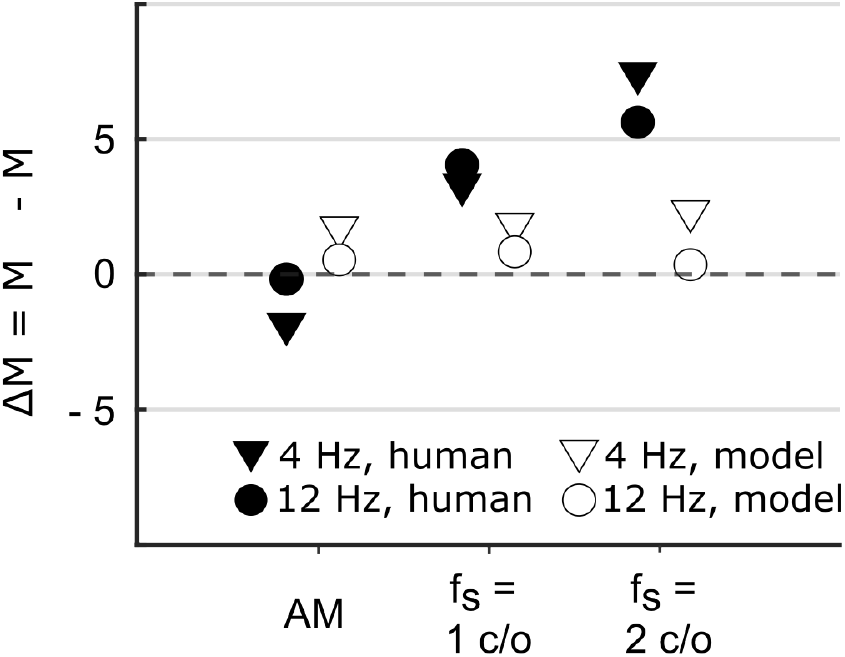
Group differences in modulation depth at threshold (ΔM = M_HI_ − M_NH_). Results are shown for human listeners (filled symbols) and CASP predictions (empty symbols) across AM and STM conditions at 4 Hz (triangle) and 12 Hz (circle). While behavioural data revealed minimal group differences for AM detection but substantial threshold elevations for HI listeners in STM conditions (up to ∼ 7.43 dB), the model predicted only modest differences between groups (up to ∼ 2.33 dB), with the largest effects observed in the 4 Hz conditions.

Figure 7 shows scatter plots of predicted versus measured thresholds for individual HI listeners across AM and STM conditions at 4 Hz (left) and 12 Hz (right). Group-averaged NH thresholds and model predictions are additionally plotted for reference. On an individual level, the model particularly fails to accurately predict thresholds from the worst performers in the STM tasks, i.e. those with the highest thresholds. Overall, when pooling across conditions, predictions correlated strongly with the behavioural data (*r* = 0.87, *p <* 0.001), indicating that CASP captured across-condition variance. However, within individual conditions, correlations did not remain significant after correction for multiple comparisons (see Table I), although it should be noted that the limited sample size (N = 10) may have reduced statistical power. Overall, the strong pooled correlation appears to be driven by the model’s ability to capture between-condition differences rather than within-condition, across-listener variability. Thus, while CASP accounts well for group-averaged performance, it fails to explain individual differences in detection thresholds.

**TABLE I.** Pearson’s correlations (*r*) and corresponding *p*-values, corrected for multiple comparisons, between measured and predicted thresholds for individual HI listeners in the different conditions.

| Condition | $r$ | $p_{corr}$ |
| --- | --- | --- |
| 4 Hz, AM | -0.36 | 0.305 |
| 4 Hz, 1 c/o | 0.20 | 0.579 |
| 4 Hz, 2 c/o | 0.64 | 0.047 |
| 12 Hz, AM | 0.34 | 0.341 |
| 12 Hz, 1 c/o | 0.61 | 0.059 |
| 12 Hz, 2 c/o | 0.10 | 0.790 |
| Overall (all conditions) | **0.87** | **< 0.001** |

**FIG. 7.**
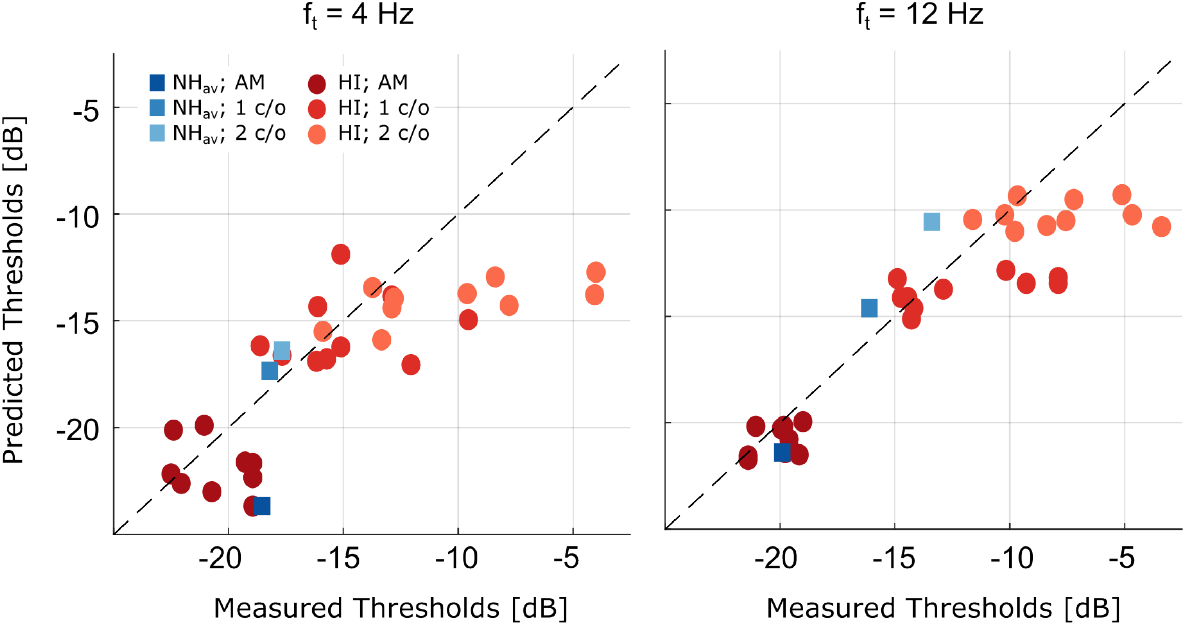
Predicted versus measured thresholds for individual HI listeners (red circles) across AM and STM conditions at 4 Hz (left) and 12 Hz (right). Group-averaged NH thresholds and model predictions (blue squares) are shown for reference. Condition types (AM, 1 c/o, 2 c/o) are indicated by shading. The dashed line indicates perfect agreement. Pooled across conditions, predictions correlated strongly with the data (*r* = 0.87, *p <* 0.001). Within-condition correlations were weaker and not significant after correction.

## V. DISCUSSION

This study evaluated the ability of the CASP model to predict STM detection thresholds in yNH and oHI listeners. The model reproduced several aspects of the measured data, including the higher sensitivity (i.e., lower thresholds) to lower temporal rates in the STM condition. However, discrepancies with the human data were also observed, particularly the inability to capture the stable performance of yNH listeners across spectral rates at the lower temporal modulation frequency, the pronounced STM deficits at higher spectral rates, and the variability across oHI listeners. The simulations provide insights into potential mechanisms contributing to STM detection in both NH and HI listeners, while also highlighting specific auditory processes that warrant further investigation.

For yNH listeners, STM detection performance was stable across spectral densities at 4 Hz but declined with increasing spectral density at 12 Hz. This pattern is consistent with earlier reports of robust low-rate modulation sensitivity and increasing difficulty when spectral and temporal modulations are combined at higher rates (Bernstein *et al*., 2013). Two mechanisms have been proposed to explain this behaviour: (i) frequency selectivity, which determines the extent of spectral smearing across channels as density increases (Bernstein *et al*., 2013; Mehraei *et al*., 2014), and (ii) TFS coding, which may provide cues for detecting slow frequency modulations at low carrier frequencies (Bernstein *et al*., 2013; Mehraei *et al*., 2014; Moore and Sek, 1996; Moore and Skrodzka, 2002). From this perspective, good performance at 4 Hz reflects reliance on TFS-based coding of spectral-peak fluctuations, whereas the rise in thresholds at higher temporal rates occurs once TFS cues are no longer available and listeners must rely on AM cues that are degraded by limited frequency resolution.

The CASP model predictions were broadly consistent with the observed NH data but offered a different mechanistic interpretation. The model reproduced the overall difference between 4 Hz and 12 Hz conditions, not through an explicit TFS mechanism, but rather through limits in modulation-phase sensitivity. At low modulation rates, residual phase cues remained accessible, supporting robust performance across spectral densities. At higher temporal rates, these phase cues were lost, and the model operated primarily on modulation power. When spectral and temporal modulations were combined, auditory filtering in the model introduces smearing across adjacent channels, reducing effective modulation depth within each channel and thereby lowering the signal-to-noise ratio at the modulation-filter outputs. In the 12 Hz conditions, the CASP predictions directly reflected this loss of within-channel modulation power, producing lower thresholds in STM compared to AM. Thus, the model suggests that limits in modulation-phase sensitivity - rather than access to carrier-level TFS - may be sufficient to explain NH performance.

The 10-Hz phase-sensitivity cutoff implemented in the model was motivated by findings from Dau (1996) who tested three normal hearing listeners’ ability to discriminate 180^°^ shifts in the starting phase of sinusoidal modulations at different modulation frequencies. Sensitivity was high at very low rates (∼ 3 Hz) but declined steeply with increasing frequency, reaching chance performance by 12 Hz. However, broader datasets on modulation-phase sensitivity remain scarce, leaving open questions about variability across listeners and the precise way to implement this within the model. Moreover, the physiological mechanisms underlying this limitation are still unidentified. Further systematic investigation is therefore needed to establish firmer constraints on this stage and to clarify its potential role in STM processing and speech perception.

Notably, the model overestimated sensitivity, i.e. predicted lower thresholds, in the 4-Hz AM condition, likely because it benefited from phase cues that human listeners may not use in simple AM detection tasks. Phase information may only be advantageous when it varies across frequency, as in the STM stimuli. The model, by contrast, treats all phase cues as informative, regardless of their distribution across channels. The failure to reproduce flat performance across spectral rates at 4 Hz therefore likely reflects an overestimation of AM sensitivity rather than an underestimation of STM thresholds. Incorporating a stage that applies modulation-phase sensitivity selectively - enhancing detection only when there is meaningful across-frequency coherence - would likely improve predictions and bring them closer to human performance.

For HI listeners, AM detection was broadly comparable to that of NH listeners, consistent with previous studies (Moore and Glasberg, 2001; Regev *et al*., 2024; Schlittenlacher and Moore, 2016; Wallaert *et al*., 2017; Wiinberg *et al*., 2019). In contrast, STM detection was substantially impaired, with threshold elevations of up to 7.43 dB relative to the NH group. CASP simulations, however, predicted only modest elevations (up to 2.33 dB) and showed little variability across individualised predictions. This mismatch suggests that the current model does not fully capture the mechanisms underlying STM deficits in HI listeners, particularly for those performing worst in the STM task. This limitation may reflect missing processing stages in the model, or shortcomings in how hearing loss is individualised. Simulated thresholds were relatively insensitive to estimates of OHC and IHC loss, with only large amounts of IHC loss producing measurable effects. This indicates that the model primarily reflected audibility limitations.

The limited influence of simulated OHC loss likely stems from the high presentation level (85 dB SPL). At such levels, the DRNL filterbank is dominated by its linear pathway, reducing frequency selectivity even in the NH configuration (Lopez-Poveda and Meddis, 2001). The OHC-loss parameter decreases nonlinear gain, effectively lowering the level at which the linear path starts to dominate. This broadens filters at low to mid levels but has little effect at higher levels, where tuning is already governed by the unchanged linear path in both NH and HI simulations. This behaviour is consistent with psychophysical findings showing minimal differences in frequency selectivity between NH and HI listeners at high presentation levels (Carney and Nelson, 1983; Dubno and Schaefer, 1991; Florentine, 1978). Low-pass filtering of the noise carrier at 2 kHz further limited the expected influence of OHC loss. The DRNL constrains the extent of implementable OHC loss at low frequencies, consistent with evidence for reduced cochlear gain in apical regions (Plack *et al*., 2008; Robles and Ruggero, 2001), making extensive OHC-related loss less likely in this range. Consequently, most of the simulated hearing loss under the present stimulus conditions was attributed to IHC dysfunction (Johannesen *et al*., 2014). This interpretation is also consistent with previous STM studies (e.g., Mehraei *et al*., 2014), which emphasised the role of frequency selectivity primarily at higher carrier frequencies not tested here. Taken together, these considerations indicate that differences in frequency selectivity between NH and HI listeners were unlikely to have substantially influenced STM detection thresholds in this study.

Instead, IHC loss emerged as the primary determinant of elevated thresholds in the simulations, although the variability introduced by this parameter remained limited. The impaired DRNL sets the operating point along the IHC nonlinearity, but the inputs reaching the IHC stage after cochlear transformation showed restricted variability under the present stimulus conditions. IHC loss is then implemented as an attenuation applied to the IHC output. This approach captures audibility limitations for large losses but appears insufficient to account for more subtle supra-threshold deficits. The precise ways in which IHC loss alters the IHC input-output function remain incompletely understood (Patra *et al*., 2024). Moreover, the IHC-loss parameter functions as a catch-all for residual threshold elevation not explained by OHC contributions, and therefore likely conflates multiple impairments rather than providing a physiologically specific description of IHC dysfunction.

The role of other possible impairments, such as possible effects of neural deafferentation, remains unclear. Existing models of transduction (Lopez-Poveda, 2014) could be integrated into the present framework to explore their potential impact on STM encoding. Importantly, modulation-phase sensitivity - critical for accounting for NH performance - may also contribute to HI deficits. In the CASP model, current OHC and IHC impairments do not alter the representation of modulation phase or the integration of cues across frequency, which may explain why STM thresholds were only minimally affected. Modelling studies of speech perception have shown that explicitly analysing coherence across auditory channels improves intelligibility predictions under phase-distorted noisy conditions (Chabot-Leclerc *et al*., 2014; Elhilali *et al*., 2003; Relaño-Iborra *et al*., 2019). However, it remains unclear how hearing loss or ageing may affect modulation-phase sensitivity or across-frequency integration. Extending CASP to include impairments in modulation-phase processing could therefore provide a mechanistic account of the pronounced STM deficits in HI listeners, but targeted empirical studies are needed to inform such model developments.

Several limitations of the present study should be considered when interpreting the findings. A first limitation concerns the use of a fixed high presentation level (85 dB SPL) without amplification for HI listeners. This choice ensured direct comparability across groups and simplified the modelling, but it may have reduced audibility at certain frequencies for some HI participants, potentially underestimating their STM sensitivity. Nevertheless, measurable thresholds were obtained for all listeners in all conditions. Elevated presentation levels can also negatively affect modulation sensitivity in NH listeners (Magits *et al*., 2019). To address these issues, Zaar *et al*. (2023) applied individualised linear amplification in STM tasks, ensuring adequate audibility while avoiding unnecessarily high levels and providing more ecologically valid listening conditions. Future work should consider adopting this approach and testing whether the model can predict STM thresholds when audibility is systematically restored. Comparisons across unaided listening, individualised linear amplification, and more realistic hearing-aid processing with nonlinear gain would provide a more stringent and ecologically relevant evaluation of the model’s predictive power.

A second limitation concerns the re-parametrisation of certain model stages that was necessary to account for the present dataset - specifically, the DRNL filterbank and the modulation phase-sensitivity stage. The revised CASP implementation (Paulick *et al*., 2025) employed relatively broad auditory filters (cf. Osses Vecchi *et al*., 2022, for model comparisons). Although this reduced frequency selectivity had little impact on simulated NH-HI group differences at the high presentation levels used here, it did affect the magnitude of the threshold reduction from AM to the 2 c/o STM condition, leading to a loss of sensitivity at 2 c/o that exceeded the empirical data. This aligns with earlier observations that limited frequency selectivity particularly hampers detection at higher spectral ripple densities, where closely spaced peaks cannot be resolved by broader filters (Mehraei *et al*., 2014). To address this, we re-instated the original DRNL implementation (Lopez-Poveda and Meddis, 2001), simplified by omitting the characteristic-frequency shift with level. These changes produced overall sharper auditory filters at high levels. In parallel, the phase-sensitivity stage was revised, as the earlier implementation produced substantial residual phase cues at higher modulation frequencies (see Appendix for implementation details). The updated stage more effectively suppressed these cues. Although these adjustments were validated for backward compatibility with prior datasets (see Appendix, Section d.), they underscore a broader concern: parameter modifications that improve descriptive accuracy for a specific dataset may compromise generalisability and complicate the functional interpretation of model parameters. Future work should therefore evaluate the robustness of these re-parametrisations across larger and more varied datasets to ensure that the model continues to reflect plausible auditory mechanisms with interpretable parameters.

Finally, a more fundamental limitation concerns the reliance on audiogram-based individualisation when attempting to predict supra-threshold performance such as STM detection. STM thresholds are thought to probe aspects of auditory processing that extend beyond audibility and are therefore only partially constrained by the audiogram. In this study, peripheral deficits were estimated in two ways: the proportion of OHC loss was derived from ACALOS loudness-growth data using a loudness model to capture recruitment, while IHC loss was inferred from the residual audiometric loss after accounting for OHC contributions. Although this partitioning provides a principled framework, it remains inherently tied to the audiogram and inherits uncertainties from the loudness-model fitting, which propagate into the individualisation process. In the present study, the impact of these uncertainties was likely limited, as STM thresholds were relatively insensitive to the precise OHC/IHC parameter values. More broadly, however, audiogram-based individualisation cannot capture other contributors to STM variability - such as synaptopathy, central auditory changes, or cognitive factors - and interpreting residual audiometric loss as IHC dysfunction risks conflating peripheral and non-peripheral sources of variability.

Still, pure-tone thresholds often covary with supra-threshold phenomena such as frequency selectivity and loudness recruitment (Sanchez-Lopez *et al*., 2020), and several studies have shown that audiogram-based predictors explain a substantial proportion of variance in speech outcomes (Bernstein *et al*., 2016, Bernstein *et al*., 2013; Zaar *et al*., 2023). Within this framework, audiogram-based models may be best viewed as defining the variance attributable to peripheral deficits, thereby clarifying the extent to which residual variability must arise from other mechanisms. This perspective also highlights the potential of STM thresholds themselves to serve as complementary individualisation metrics, extending beyond audiogram-based parameters and enabling more comprehensive accounts of inter-individual differences in speech perception.

## VI. CONCLUSION

This study examined STM detection in NH and HI listeners and evaluated the ability of an individualised auditory model to predict these STM detection thresholds. CASP reproduced general threshold patterns across temporal and spectral rates but failed to fully capture group differences and the pronounced variability across individuals. Model predictions showed limited sensitivity to outer and inner hair cell loss estimates, indicating either that other mechanisms contribute to STM deficits or that current implementations of these impairments are insufficient. Within the model framework, modulation-phase sensitivity emerged as a key factor for explaining NH performance at low and high temporal rates, highlighting its potential importance for understanding STM processing more broadly. Future work should investigate the role of modulation-phase sensitivity in speech perception and examine how hearing loss alters this mechanism, alongside contributions from other mechanisms, such as neural deafferentation, central auditory changes, and age-related and cognitive factors. Moreover, future work could explore incorporating STM thresholds as an individualisation metric, beyond traditional audiogram-based approaches. This may enable the development of auditory models that more accurately predict speech performance and provide a stronger basis for hearing-aid evaluation.

## ACKNOWLEDGMENTS

We would like to thank Jonathan Regev for valuable help with the experimental design and setup and providing code for the ACALOS experiment, as well as Johannes Zaar for providing base code for generating the STM stimuli. This work was carried out in connection to the Center for Applied Hearing Research (CAHR) supported by Widex, Oticon, GN ReSound, and the Technical University of Denmark.

## DATA AVAILABILITY

Data and model implementation will be made available upon request.

## APPENDIX A: APPENDIX: MODEL ADJUSTMENTS

Select adjustments were made to the model implementation compared to that described in Paulick *et al*. (2025). The following sections provide a detailed account of these changes, followed by an assessment of their backward compatibility. Specifically, the modified model was re-evaluated on the same benchmark experiments used in Paulick *et al*. (2025) - intensity discrimination, forward masking, and modulation detection - to verify that its predictive performance was maintained.

### 1. DRNL filterbank

For the NH simulations, the original parametrisation of the DRNL model proposed by Lopez-Poveda and Meddis (2001) was adopted, rather than the implementation used in Paulick *et al*. (2025). The key difference lies in the number of cascaded gammatone and low-pass filters in the linear and nonlinear paths. In the modified linear path, three gammatone filters and four low-pass filters were applied (compared to two gammatone and four low-pass filters in Paulick *et al*. (2025)), while the nonlinear path comprised a cascade of three gammatone and three low-pass filters (compared to two and one, respectively, in Paulick *et al*. (2025)). The DRNL implementation of Paulick *et al*. (2025) was aligned with that used in the speech-based sCASP model (Relaño-Iborra *et al*., 2019). A com-parative study of auditory models demonstrated that this version produced comparatively broad filters (Osses Vecchi *et al*., 2022). By reverting to the original DRNL (Lopez-Poveda and Meddis, 2001) parametrisation, we aimed to restore sharper frequency selectivity which proved necessary to capture the trends in the STM data. In addition, one simplification was introduced: the centre frequencies and cut-offs of the gammatone and low-pass filters were fixed to the characteristic frequency, independent of level. Unlike the original formulation, no level-dependent frequency shifts with increasing sound pressure level were implemented here. This choice reflects more recent findings that challenge the notion of characteristic-frequency shifts at higher levels (Moore and Glasberg, 2003).

### 2. Modulation phase sensitivity

The implementation of the modulation phase-sensitivity stage was slightly modified compared to earlier work, while the underlying rationale remained unchanged — namely, to reduce phase information in the internal representation above 10 Hz. In the present study, the real output of the modulation filterbank was first obtained. For filters with centre frequencies at and above 10 Hz, the Hilbert envelope of this real output was then calculated and subsequently low-pass filtered with a 10 Hz cutoff, while preserving the overall energy at the filterbank output. For filters centred below 10 Hz, the real filter output was used directly. This procedure produced a stronger suppression of phase information at high modulation frequencies than the earlier implementation. In previous work, phase sensitivity had instead been reduced by taking the real part of the modulation filter output for *f*_mod_ ≤ 10 Hz and the absolute value for *f*_mod_ *>* 10 Hz.

### 3. Back-end stage

In the model backend, a supra-threshold template was generated prior to each run and correlated with the three alternative intervals across all dimensions of the internal representations. The correlation was computed as:

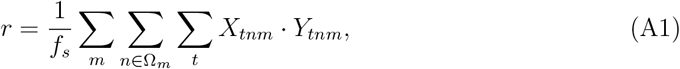

where *f*_*s*_ is the sampling frequency, *X* = [*T* × *N* × *M*] is the interval signal, and *Y* = [*T* × *N* × *M*] is the template. The indices *t, n*, and *m* refer to time, auditory channel, and modulation channel, respectively, while Ω_*m*_ denotes the subset of auditory channels selected by the channel-selection algorithm of Paulick *et al*. (2025), and constrained such that *f*_*n*_ *>* 4*f*_*m*_ for each modulation frequency *f*_*m*_. This non-normalised correlation approach differs slightly from Paulick *et al*. (2025), where the average over time and frequency was subtracted from both the signal and the template before computing the correlation. Due to the re-implementation of the modulation phase sensitivity for modulation frequencies above 10 Hz, where essentially only the DC component (modulation power) remains, a normalised correlation approach was not feasible.

### 4. Backwards compatibility

The backward compatibility of these adjustments was evaluated by rerunning the benchmark experiments reported in Paulick *et al*. (2025), including intensity discrimination, forward masking, and modulation detection for NH listeners. Table II presents the mean absolute error and Pearson correlation with human data for each task, comparing the predictions of the previous model with those obtained using the modified implementation described above. Overall, similar performance of the two model implementations can be observed across the different experimental conditions.

**TABLE II.** Backward compatibility assessment: Mean absolute error (MAE) and Pearson correlation (*r*) comparing the previous CASP implementation (Paulick *et al*., 2025) and the modified implementation across benchmark experiments.

| Experiment | Condition | MAE | | Pearson $r$ | |
| --- | --- | --- | --- | --- | --- |
|  |  | Paulick 2025 | proposed | Paulick 2025 | proposed |
| Intensity Discrimination | Tone | 0.27 dB | 0.20 dB | 0.807 | 0.835 |
| Intensity Discrimination | Broadband noise | 0.67 dB | 0.10 dB | 0.224 | -0.334 |
| Forward Masking | 40 dB | 3.47 dB | 5.00 dB | 0.984 | 0.992 |
| Forward Masking | 60 dB | 2.24 dB | 5.25 dB | 0.991 | 0.999 |
| Forward Masking | 80 dB | 1.70 dB | 4.94 dB | 0.996 | 0.995 |
| Modulation Detection | 3 Hz | 6.14 dB | 8.79 dB | 0.981 | 0.939 |
| Modulation Detection | 31 Hz | 3.42 dB | 2.02 dB | 0.798 | 0.846 |
| Modulation Detection | 314 Hz | 2.21 dB | 1.65 dB | 0.495 | 0.953 |

